# msCRUSH: fast tandem mass spectra clustering using locality sensitive hashing

**DOI:** 10.1101/308627

**Authors:** Lei Wang, Sujun Li, Haixu Tang

## Abstract

Large-scale proteomics projects often generate massive and highly redundant tandem mass (MS/MS) spectra. Spectra clustering algorithms can reduce the redundancy in these datasets, and thus speed up the database searching for peptide identification, a major bottleneck for proteomic data analysis. Furthermore, the *consensus spectra* derived from highly similar MS/MS spectra in the same cluster may enhance the signal peaks while reduce the noise peaks, and thus will improve the sensitivity of peptide identification. In this paper, we present the software msCRUSH, which implemented a novel spectra clustering algorithm based on the locality sensitive hashing (LSH) technique. When tested on a large-scale proteomic dataset consisting of 18.4 million spectra (including 11.5 million spectra of charge 2+), msCRUSH runs 7.6-12.1x faster than the state-of-the-art spectra clustering software, PRIDE Cluster, while achieves higher clustering sensitivity and comparable accuracy. Using the consensus spectra reported by msCRUSH, commonly used spectra search engines MSGF+ and Mascot can identify 5% and 4% more unique peptides, respectively, comparing to the identification results from the raw MS/MS spectra at the same false discovery rate (1% FDR) of peptides. msCRUSH is implemented in C++, and is released as open source software.

## Introduction

With the rapid technical advancement in the past decade, liquid chromatography coupled tandem mass spectrometry (LC-MS/MS) has achieved significantly higher throughput and sensitivity, and thus extensive applications in large-scale analyses of complex proteomic samples.^1-3^ The current mass spectrometry instruments normally acquire 10^4^-10^5^ MS/MS spectra on proteome samples within a few hours in a single run, from which tens of thousands of peptides can be identified.^3^ To further improve sensitivity, analytical and/or biochemical approaches can be utilized to fractionate proteomic samples to reduce their complexity prior to LC-MS/MS analyses. For instance, reversed-phase peptide fractionation combined with LC-MS/MS, often referred to as the *shotgun proteomics* approach,^4^ leads to high resolution peptide separation, resulting in the identification of a half million unique peptides from over 14,000 protein isoforms in a single human cell line.^1^ Biochemical fractionation approaches, e.g., based on subcellular compartments, ^5^ can substantially increase the identification coverage of the target proteome, generating even more massive mass spectra datasets. Finally, large-scale proteomic projects (e.g., the CPTAC project^6^) acquire MS/MS data from hundreds to thousands of samples (e.g., collected from patients and healthy individuals), each of which is analyzed with multiple technical replicates. As a result, these projects typically produce datasets consisting of over 10^8^-10^9^ MS/MS spectra.

The whole set of spectra acquired in a large scale proteomics project is often highly redundant, i.e., many spectra are from the same peptides, and thus exhibit high similarity.^7^ There are various reasons causing this redundancy: 1) in a single LC-MS/MS analysis, the MS/MS may isolate and fragment the same (abundant) peptide ion multiple times (depending upon sample complexity and MS instrument setting); 2) when samples are fractionated, some peptides may be present in multiple fractions; and 3) a majority of peptides may be present in multiple samples analyzed in the single project. Searching redundant spectra from highly abundant peptides could be both time and resource consuming. Notably, such redundancy in a large set of spectra can be significantly reduced prior to peptide identification using database search engines (e.g., Mascot,^8^ Sequest^9^ or MSGF+^10^) by clustering MS/MS spectra based on their similarities such that the spectra in the same cluster are likely generated from the same peptide and then retaining only a representative spectrum in each cluster for subsequent database searching. This spectra clustering approach was commonly adopted for building an MS/MS spectra library,^11^ in which a *consensus* spectrum was used as the representative for each cluster. There are several advantages to exploit this approach in proteomic data analyses. First, after spectra clustering, much fewer consensus spectra (comparing to the original set of spectra) need to be identified by database searching, and thus the processing time for peptide identification will be significantly reduced. Second, the consensus spectra often show high signal-to-noise ratio, and are more likely to be accurately identified. Finally, the clusters consisting of large number of spectra, even if their consensus spectra are not identified, are worth further investigation because they are often generated from abundant, novel molecules, e.g., the peptides containing mutations or post-translational modifications (PTMs).

Conventional mass spectrum clustering algorithms, e.g., MS-Clustering,^12^ MaRaCluster,^13^ and PRIDE Cluster,^14^ adopted the one-against-all approach, whereas each spectrum is compared against other spectra of the same charge and within a precursor mass precision using various similarity measures such as cosine similarity^7^ and self-defined metrics.^13-15^ Given a certain similarity threshold, the pair of spectra with highest similarity, higher than the given threshold, will be grouped into a single cluster. A consensus spectrum is then constructed from the newly formed cluster to replace the two similar spectra. The procedure terminates if no pair of spectra remains with similarity higher than the given threshold. For large mass spectra datasets (e.g., containing millions of spectra), this algorithm is slow because the pairwise comparison involves each pair of spectra of the same charge and close precursor mass. To speed up the algorithm for clustering large spectra set, heuristics were developed to avoid pairwise comparisons. For instance, MS-Clustering^12^ considers only pairs of spectra that share at least one peak among their top five strongest peaks, respectively. Both MS-Clustering and PRIDE Cluster^14^ adopt a greedy clustering strategy where they merge first pair of spectra with similarity over the given threshold instead of merging the pair of spectra with the highest similarity, in each round of clustering with decaying similarity threshold.

In this paper, we present a novel spectrum clustering algorithm based on the locality sensitive hashing (LSH) technique. We showed that our algorithm can significantly speed up clustering by selecting a subset of highly similar spectra through one-time processing of all spectra while retaining comparable or higher sensitivity and accuracy. We implemented this algorithm in a software package named msCRUSH (standing for <u>m</u>ass <u>s</u>pectrum <u>C</u>luste<u>R</u>ing <u>U</u>sing locality <u>S</u>ensitive <u>H</u>ashing) in C/C++. As demonstrated on a large-scale human proteomic dataset consisting of 18.4 million spectra (including 11.5 million spectra of charge 2+), it runs 7.6-12.1x faster than the state-of-the-art spectra clustering software PRIDE Cluster,^14^ while our algorithm achieved higher clustering sensitivity and comparable accuracy. We also showed that, using the consensus spectra for the spectrum clusters reported by msCRUSH, about 5% or 4% more unique peptides could be identified when MSGF+^10^ or Mascot^8^ was used as the database search engine, respectively, comparing to the identification results from the raw un-clustered spectra, and about 17% or 7% more unique peptides comparing to the results from using the consensus spectra reported by PRIDE Cluster. Consensus spectra generated by msCRUSH can also speed up database searching compared to un-clustered original spectra, regardless of peptide search engines: 2.37x is achieved using MSGF+ and 2.23x is achieved using Mascot, both on a large-scale dataset consisting of 11.5 million spectra of charge 2+.

## Materials and Methods

The iterative clustering algorithm implemented in msCRUSH is illustrated in Figure 1. Instead of computing spectral similarity for each pair of spectra, which is computationally intensive, we adopt the locality sensitivity hashing (LSH) technique to group similar spectra into *buckets* in a LSH table, and then compute the pairwise similarities between spectra within each bucket that share the same charge and close precursor mass. The property of LSH ensures that two similar spectra are likely to be grouped into the same bucket. Performing pairwise similarity computations in each bucket that contains a small set of potentially similar spectra substantially reduces the computational cost. In addition, to increase the sensitivity of the approximate LSH technique (in particular when the bucket size is large), our algorithm uses gradually decaying similarity thresholds, starting from stringent to loose, during the iterative process. Below, we describe the iterative clustering algorithm in details.

**Figure 1:**
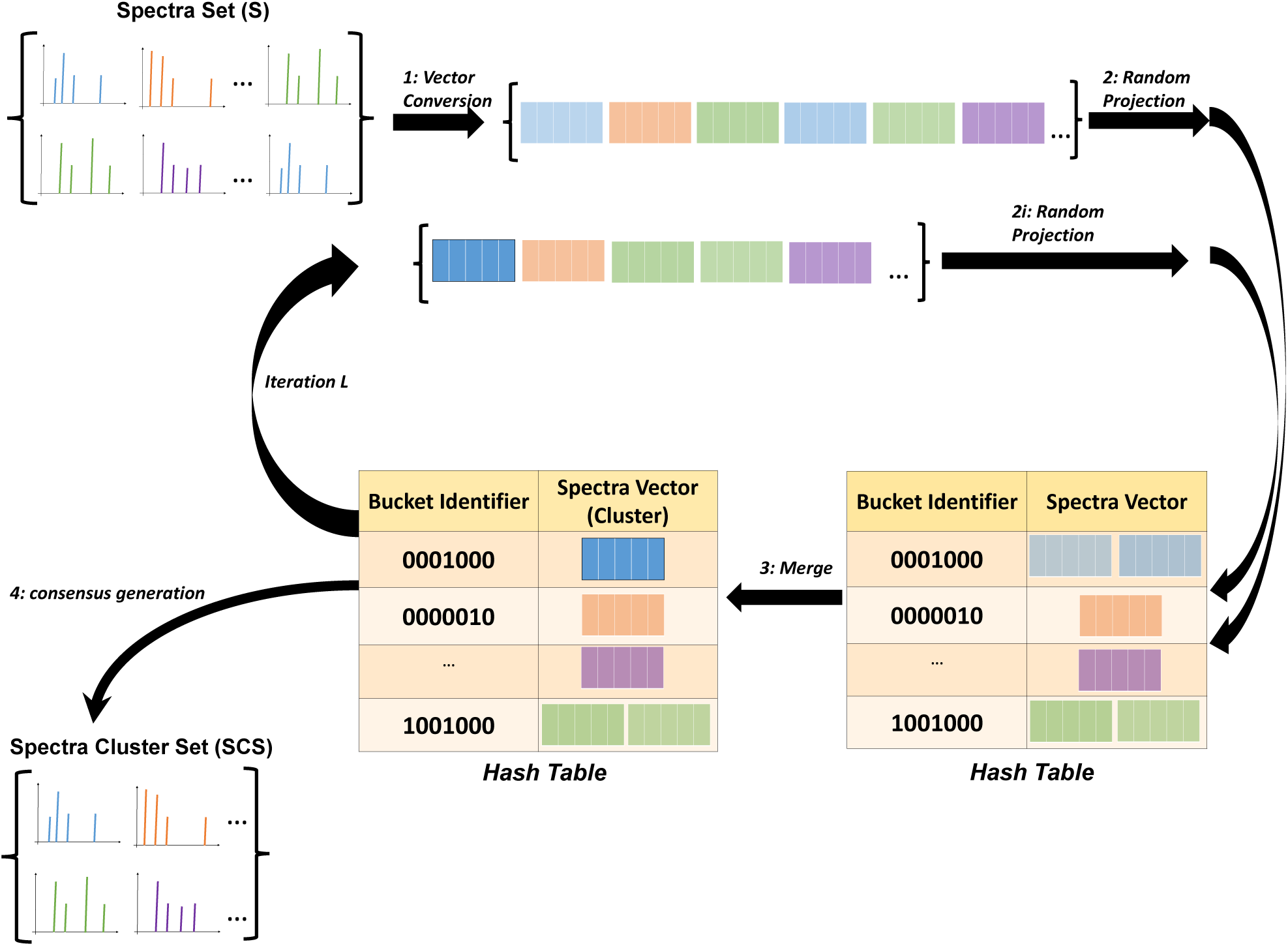
The workflow of msCRUSH algorithm. 1: Vector conversion: Prior to spectrum clustering, each input MS/MS spectrum is first pre-processed and embedded into a numerical vector. 2: Random projection: those vectors are then randomly projected into buckets by utilizing the selected SimHash functions. 3: Merge: Within each bucket, cosine similarity between each pair of spectra of the same charge and close precursor mass is calculated. If the similarity is higher than the specific threshold, they will be merged into a consensus spectrum; otherwise, they will remain separate. Iteration L: After merging and replacement, the new spectrum (i.e. consensus spectrum) vector will proceed into next iteration of vector conversion, random projection and merge. 4: Consensus generation: After a maximum number of iterations, msCRUSH will generate consensus spectra as the final clustering report.

### Measuring similarity between two tandem mass spectra

We consider the tandem mass (MS/MS) spectra clustering problem as a high dimension vector clustering problem: each input MS/MS spectrum consisting of hundreds of peaks is considered as a *sparse* vector, in which each dimension is represented by a bin of mass-to-charge (m/z) ratio, with the bin size representing the resolution of the mass spectrometer. The normalized peak intensity is assigned to the vector corresponding to the m/z. The vector clustering problem has been extensively studied in many applications including bioinformatics, such as for the clustering of gene expression profiles^16,17^ and text documents.^18-20^ Similar to the other applications, the similarity between two vectors (in our case, MS/MS spectra) can be measured by different metrics. In this paper, we used the *cosine similarity*, a variant of angle-based distance^7^ between the vector representations of MS/MS spectra, which can be approximated using the LSH technique. The high cosine similarity between two MS/MS spectra indicates that they are likely to result from the same peptide. The other similarity measures, including the Pearson correlation coefficient ^21^ and the Euclidean distance ^22^ have been shown to be highly correlated with the cosine similarity.^23^

### Locality Sensitive Hashing (LSH) for spectra similarity search

Searching similar vectors based on cosine similarity can be accelerated by using locality sensitive hashing (LSH) algorithms as proposed previously.^24,25^ Notably, in computational proteomics, an LSH technique called MinHash was adopted to speed up database searching for peptide identification.^23^ However, MinHash approximates the size of intersection set between two sets of fragment peaks (e.g., one from an experimental spectrum and the other from the theoretical spectrum of a peptide in the database), and thus is not appropriate for approximating the similarity between two vectors. In a brief summary, LSH utilizes hash function *y* = *h*(*x*), where *y* is the hash value, to map objects into *buckets* in a hash table, which can be indexed by their hash values (*codes*). Conventional hash functions are designed to map objects to different buckets if these objects are not exactly the same, thus to reduce the chance of collisions. In contrast, LSH functions attempt to assign similar objects (in our case, MS/MS spectra) into the same buckets with higher probabilities than objects with low similarity.

Generally, LSH is defined in following: Let 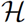 be a family of hash functions *h* that maps objects in a metric space *M* to a bucket *s ∈ S*. The family 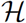 is called *locality sensitive* under a threshold *R*, an approximation factor *c (c >* 1) and collsion probability *P*_1_ and *P_2_ (P_1_* > *P*_2_), if for any two objects *p, q ∈ M*,

- if *d(p,q) ≤* R, then *Pr*[h(p) = h(q)] ≥ *P*_1_,
- if *d(p,q) ≥* cR, then *Pr*[h(p) = h(q)] ≤ *P*_2_.

where *d(p, q)* is the distance between objects *p* and *q* (where *p* and *q* are vectors, *d(p, q)* = ||*p* − *q*||), and *Pr*[*h*(*p*) = *h*(*q*)] represents the probability that *p* and *q* collide under a hash function. Specifically, in this paper, we used a family of LSH functions called SimHash,^24,25^ which aims to approximate the cosine similarity measure (i.e., the angle between two vectors *s_i_* and 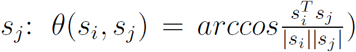 using random projection,^26^ when searching for similar spectra. Given two vectors *s_i_* and *s_j_*, the SimHash function is defined as *h*(*x*) = *sign(w^T^x)*, where w is a randomly chosen vector and *x* is the input item; *h*(*x*) = +1 or −1 depending on which side of the hyperplane (defined by *w*) *x* lies. For any *s_i_* and *s_j_*, 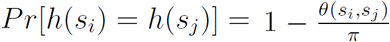. Hence, the higher similarity (i.e., smaller *θ*) between the two vectors, the more probable they are mapped to the same bucket in a hash table by the SimHash function *h*.

### Iterative clustering

Previous spectra clustering algorithms such as MS-Clustering^12^ and PRIDE Cluster^14^ follow the *one-against-all* strategy. These methods attempt to measure the similarity between all pairs of spectra of the same charge and close precursor mass (within certain mass precision). The general spectra clustering algorithm can be considered as a special case of the hierarchical agglomerative clustering (HAC)^27^, which starts from the clusters each consisting of one spectrum (i.e., a *singleton*), and in successive steps merges two clusters into a single cluster if they represent spectra of the same charge, close precursor mass and high spectra similarity. Rigorous HAC algorithms run in time complexity of 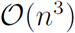 for *n* items (spectra), while it can be reduced to 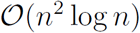 at the cost of using more memory.^27^ Due to the large number (e.g., 11.5 million charge 2+ spectra in the test dataset) of MS/MS spectra, even the optimized hierarchical clustering scheme runs very slow in practice. To address this issue, both MS-Clustering and PRIDE Cluster adopted an iterative, greedy strategy to obtain approximate hierarchical clusters: it merges an *arbitrary* pair of spectra with similarity above a given threshold instead of merging two most similar spectra^12,14^ in each round of clustering. Only as few as 4 rounds of clustering were performed by default in PRIDE Cluster for the sake of running time. As a result, these algorithms did not enumerate all pairs of spectra with the same charge and close precursor mass in each step, and thus may sacrifice some accuracy, resulting in both false positives (i.e., spectra of different peptides that were grouped into the same cluster) and false negatives (i.e., spectra of the same peptides that were assigned to various clusters).

In an attempt to improve the clustering accuracy at the minimum computational cost, we adopted an iterative decay approach as well. Instead of performing pairwise computations among spectra of the same charge and close precursor mass and then merging pairs of spectra with similarity exceeding a given threshold^12,14^ that is particularly computationally expensive for a set of large number of spectra, msCRUSH only computes the pairwise similarity of a set of highly similar spectra residing in each hash bucket. As shown in Figure 1, while we conduct multiple rounds of SimHash-based clustering, we decrease the similarity threshold in the successive iterations. For instance, the similarity threshold is set to be stringent (e.g., 0.9) for the first round (used interchangeably with iteration), and with the clustering proceeding to the next rounds, we continue decreasing the threshold (e.g., to 0.85 and so on) until it reaches a preset minimum value (e.g., 0.65).

### Augmenting LSH functions

To optimize the LSH-based clustering algorithm, we need to adjust two embedded parameters: the number of hash functions in one hash table and the number of hash tables (in our case, iterations), as shown in Figure 2(a) and 2(b), respectively. When multiple hash functions are used, the LSH algorithm amplified the gap of the collision probability between the similar spectra and the dissimilar spectra. In particular, a *compound* hash function *g*(*x*) = (*h*_1_(*x*),…, *h_K_*(*x*)) is constructed by concatenating *K* SimHash functions, *h*_1_(*x*),…,*h_K_*(*x*), where each SimHash function *h_K_*(*x*) is chosen randomly from the family 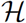.^26^ Two spectra are mapped into the same bucket in a hash table of *K* concatenated SimHash functions, only if their *compound* hash keys are same; hence for two spectra with a collision probability *p* under a single SimHash function h, their collision probability under the *compound* hash function becomes *p^K^*. As shown in Figure 2(a), using only one hash function (*k* =1, the blue line), the collision probability (y-axis) for two spectra to be mapped into the same bucket in a single hash table is linearly correlated with their cosine similarity (x-axis). When multiple hash functions are used (e.g., *k* = 10, the red line), the spectra similarity and collision probability show nonlinear correlations. For example, when 10 hash functions are used, two similar spectra with cosine similarity of 0.7 is much more likely to be mapped into the same bucket than two dissimilar spectra with similarity 0.2: the collision probability is 0.7^10^ ≈ 0.03 for the two similar spectra, whereas the probability is 0.2^10^ ≈ 10^−8^ for the two dissimilar spectra. However, using multiple hash functions also decreases the collision probability of two similar spectra, and thus increases the number of false negatives in clustering because two similar spectra are likely mapped to different buckets in the compound hash table.

**Figure 2:**
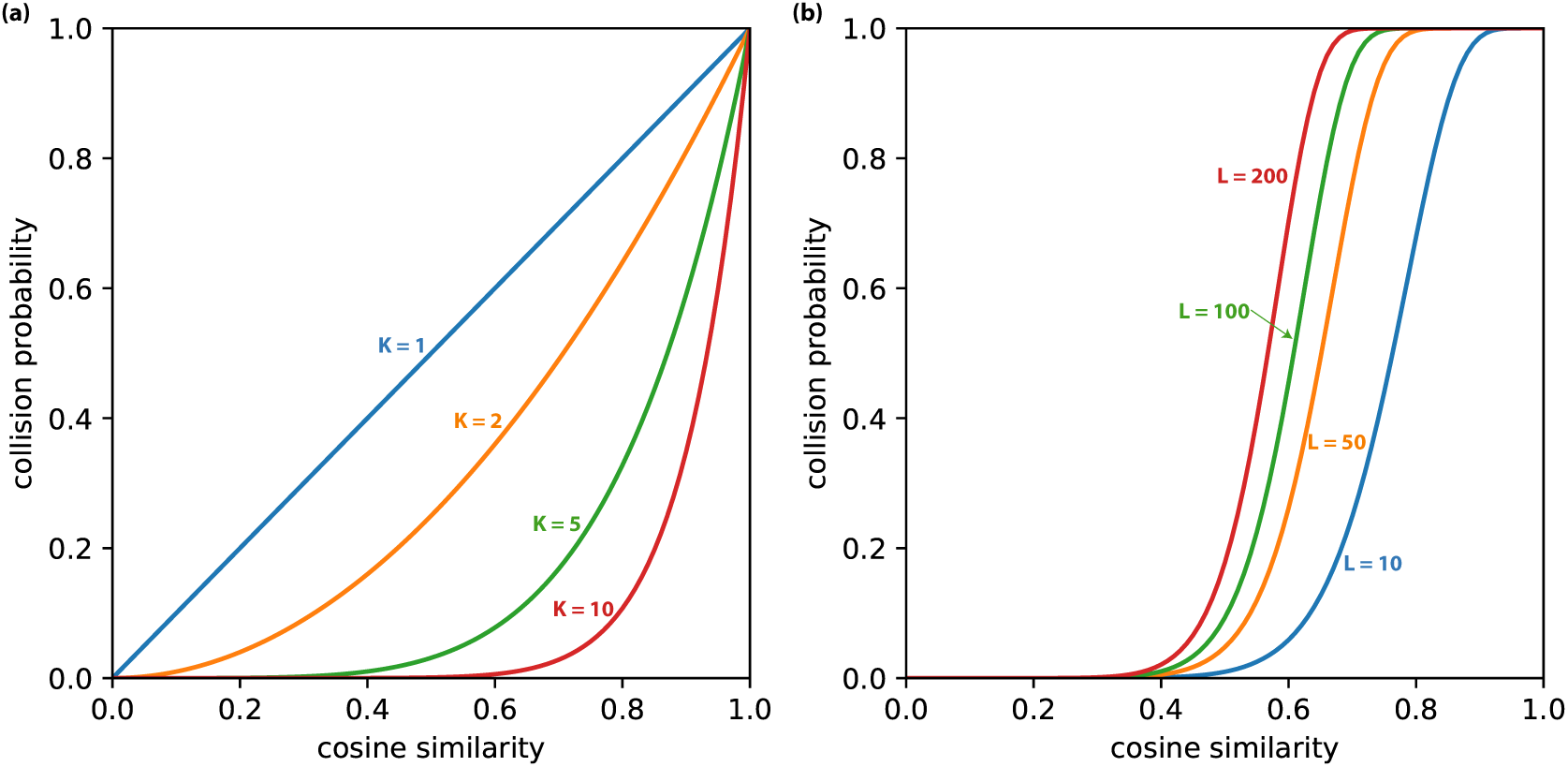
Illustration of the collision probability between two spectra under various compound SimHash functions and various hash tables. (a) The collision probability decreases as more hash functions are concatenated in a compound hash function, i.e. a hash table.; (b) Augmented SimHash functions (10 hash functions are concatenated per hash table) incorporating multiple hash tables can increase the collision probability for pairs of similar spectra, while keeping the collision probability between dissimilar spectra relatively low.

To address this issue, we performed multiple iterations of clustering, and a hash table consisting of one compound hash function is selected in each iteration. We then attempted to verify the similarity of two spectra if they are mapped to the same bucket from at least one of *L* hash tables (i.e. from at least one iteration). In this case, the collision probability of two spectra with similarity *p* becomes 1 − (1 − *p^K^*)*^L^*. For *k* = 10 and *L* = 100 (Figure 2(b), green line), the collision probability is ≈ 0.94 for two similar spectra with cosine similarity 0.7, whereas ≈ 10^−5^ for two dissimilar spectra with cosine similarity 0.2. For each iteration, the spectra within the same bucket that share the same charge and similar precursor mass are retrieved as putative similar spectra, on which the similarities are computed explicitly. If the similarity is higher than the threshold for the specific iteration, they will be merged into a consensus spectrum (see below for details). In each iteration, we decrease the similarity threshold based on a decay function: 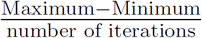, where the maximum and minimum number similarity are set as 0.90 and 0.65, respectively. For instance, for a total of 100 iterations, each iteration will decrease the threshold by 0.0025. Overall, during multiple iterations, the LSH-based clustering algorithm can significantly reduce the running time for computing pairwise similarity between spectra (as dissimilar spectra are not likely mapped to the same buckets in each iteration and thus expensive pairwise similarity computations between dissimilar spectra can be avoided) while retaining high sensitivity for grouping similar spectra (as similar spectra are likely to be mapped to the same bucket in at least one iteration).

### Generating consensus spectra

A *consensus spectrum* is the representative mass spectrum for the (similar) spectra in each cluster. Previous work^12,28^ showed that representing a cluster of highly similar spectra by a consensus spectrum can improve the signal-to-noise ratio of the mass spectrum. In msCRUSH, initially, each cluster consists of only one spectrum, which will be the consensus spectrum for the cluster. For each successive iteration, if the similarity of two colliding (consensus) spectra is above the threshold, the corresponding clusters are merged into a single cluster, simultaneously generating a new consensus spectrum to represent the whole cluster of spectra. Subsequently, the cosine similarity is computed between the resulting consensus spectrum and the other consensus spectra, each representing a remaining cluster. This simple process avoids the expensive computation of similarity between each pair of spectra in two clusters. Note that the number of consensus spectra always decreases after each iteration, and thus the final output of the clustering algorithm is a set of consensus spectra, which are much fewer than the set of raw spectra. These consensus spectra can be used as input for database searching to reduce the computing time for peptide identification.

Given a cluster of MS/MS spectra, the consensus spectrum is computed by merging the peaks in all input spectra: the peaks in all spectra within certain mass precision are merged into a single peak in the consensus spectrum, for which the m/z is set as the average m/z of all merged peaks weighted by the intensity of each peak, and the intensity is set to be the average intensity of all merged peaks. Intuitively, the consensus spectrum representing *N* MS/MS spectra in a cluster has a high signal-to-noise ratio, because the intensity of a noise peak (occurring in only one input spectrum) is reduced by *N* times while the the intensity of a signal peak (occurring in all input spectra) remains about the same.

### Spectrum pre-processing

Prior to spectrum clustering, each input MS/MS spectrum is first pre-processed to remove putative noise peaks. Specifically, we partition the peaks in the spectrum into 100-Da wide bins (from 200 Da to 2,000 Da), and in each bin, only the five strongest peaks are retained. In order to reduce the influence of dominant peaks on the cosine similarity, the natural logarithm of each peak’s intensity is considered in spectra clustering. Finally, the intensities of all peaks are normalized such that the intensity of the strongest peak in a spectrum is set to be the same (1,000 by default in the msCRUSH implementation).

### MS/MS datasets

We used two benchmark datasets of MS/MS spectra for testing the performance of msCRUSH and PRIDE Cluster in this paper. Dataset A from ProteomeXchange^29^ (ID:PXD001197) contains 903,237 spectra in total, including charges 2+, 3+ and 4+. This dataset was acquired using the high resolution mass spectrometry LTQ Orbitrap Elite on the human cell line HEK293.^30^ Dataset B from ProteomeXchange(ID: PXD000561) contains 18,405,828 spectra in total, including charges 1+, 2+, 3+ and 4+. Detailed numbers of spectra with each charge state for these datasets can be found in Table 1. The MS/MS spectra were acquired in a comprehensive human proteomic study using high-resolution Fourier-transform mass spectrometry, while analyzed 30 histologically normal human samples, including 17 adult tissues, 7 fetal tissues and 6 purified primary haematopoietic cells.^31^

**Table 1:** Comparison of msCRUSH and PRIDE Cluster algorithms. Running time was measured in seconds using 40 threads for both msCRUSH and PRIDE Cluster. The results were compared in terms of purity, within-cluster entropy (WCE) and within-peptide entropy (WPE). The number of spectra was calculated in thousands. The cosine similarity threshold for merging spectra was set to 0.65 for each dataset, and 15 hash functions, 100 iterations were used in msCRUSH.

| Index | No. of spectra | No. of Clusters |  | Time for Clustering |  | Purity |  | WCE |  | WPE |  |
| --- | --- | --- | --- | --- | --- | --- | --- | --- | --- | --- | --- |
| Dataset |  | PRIDE(%) | msCRUSH(%) | PRIDE | msCRUSH | PRIDE | msCRUSH | PRIDE | msCRUSH | PRIDE | msCRUSH |
| A-2+ | 415 | 261(62.89) | 148(35.66) | 237 | 75 | 0.992 | 0.978 | 0.022 | 0.068 | 2.041 | 0.747 |
| A-3+ | 358 | 292(81.56) | 217(60.61) | 225 | 67 | 0.996 | 0.991 | 0.012 | 0.027 | 2.779 | 1.248 |
| A-4+ | 111 | 98(88.29) | 85(76.58) | 83 | 19 | 0.999 | 0.998 | 0.002 | 0.005 | 2.772 | 1.680 |
| B-1+ | 762 | 677(88.85) | 557(73.10) | 377 | 32 | 0.996 | 0.993 | 0.011 | 0.021 | 2.234 | 0.538 |
| B-2+ | 11,525 | 7,695(66.77) | 5,623(48.79) | 9,480 | 783 | 0.989 | 0.980 | 0.032 | 0.070 | 2.912 | 1.835 |
| B-3+ | 4,981 | 3,714(74.56) | 3,216(64.57) | 2,812 | 301 | 0.995 | 0.993 | 0.014 | 0.022 | 3.231 | 2.121 |
| B-4+ | 1,138 | 942(82.78) | 874(76.80) | 604 | 80 | 0.997 | 0.997 | 0.009 | 0.011 | 3.179 | 2.569 |

### Database Searching

We used MSGF+^10^ and Mascot^8^ database search engines for peptide identification. The parameters for database searching is set as the following to match the experimental conditions:^30,31^ 1) Instrument type: high-resolution LTQ; 2) Precursor mass tolerance: 50ppm; 3) Isotope error range: −1, 2; 4) Modification: oxidation as variable and carboamidomethy as fixed; 5) Maximum charge: 7; and 6) Minimum charge: 1. The false discovery rate (FDR) is estimated by using a target-decoy search (TDA) approach,^32^ in which the decoy proteins were generated by reversing the protein sequences in the target Uniprot human database. ^33^

### Evaluation Metrics

A variety of metrics can be employed to evaluate the performance of spectra clustering algorithms. We evaluated both the running time and the quality of the spectra clustering algorithms of msCRUSH and PRIDE Cluster. To evaluate the quality of clustering algorithms, we adopted several metrics defined by Enrique *et al*,^34^ including cluster homogeneity (purity), cluster completeness (within-cluster entropy), and peptide completeness (within-peptide entropy), each of which focuses on one aspect of the quality of clusters. We describe these metrics in details below. To be noted, in a typical LC-MS/MS experiment, on average, around 25% spectra can be identified at a low false discovery rate (1%).^14^ Our evaluation utilizes these identified spectra as golden standard to compute the evaluation metrics. In reality, because a small fraction of these spectra are not correctly identified, the metrics may not be completely accurate.

**Purity**. Cluster precision (CP) is defined as the frequency of the most commonly identified peptide (*p*) by the spectra in each cluster. Purity is then computed as the average of cluster precision on all clusters weighted by the cluster size.

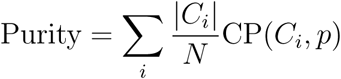

where CP is the cluster precision and *N* is the total number of identified spectra, |*C_i_*| is the number of identified spectra in cluster *C_i_*. A high purity (close to 1.0) indicates the clusters are mostly homogeneous, and thus have high sensitivity.

**Within-cluster entropy**. Within-cluster entropy (WCE) indicates how diversified the identified peptides are within a cluster,

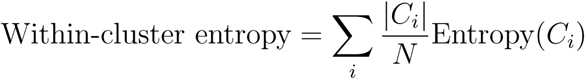

where *N* represents the number of identified spectra, |*C_i_*| is the number of identified spectra in the cluster *C_i_* and Entropy(*C_i_*) represents the entropy of the identified peptides in the cluster *C_i_*, which is defined as

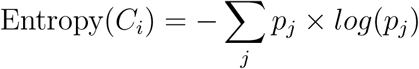

where *p_j_* is the frequency of the peptide *j* identified by the spectra in cluster *C_i_*. The within-cluster entropy is 0 if this cluster contains spectra all identified as the same peptide. A high within-cluster entropy indicates a low sensitivity of the clustering algorithm, as a variety of peptides are grouped into the same cluster.

**Within-peptide entropy**. The within-peptide entropy (WPE) measures how spectra of the same peptides distribute over diverse clusters,

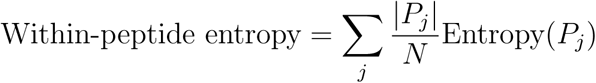

where *N* is the total number of identified spectra, | *P_j_* | is the number of spectra identified from the same peptide *P_j_* across all clusters, and Entropy(*P_j_*) represents the entropy of the peptide *P_j_*, which is defined as

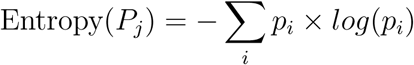

where *p_i_* is the frequency of spectra identified as the peptide *P_j_* from cluster *C_i_*. A high within-peptide entropy indicates a low sensitivity of the clustering algorithm, because all spectra from the same peptides are not grouped into the same cluster.

### Software availability

We implemented our algorithm including the pre-processing and the clustering algorithm in C++ in the software package msCRUSH, which is portable across different platforms, e.g., Windows, Linux, Unix and Mactonish. The functionality of multi-threading was implemented in msCRUSH using OpenMP ^35^ to further speed up the spectra clustering. msCRUSH is released as open-source software at github: https://github.com/COL-IU/msCRUSH

## Results

### Default parameters

The performance of msCRUSH depends upon several parameters: the number of hash functions in one hash table, the number of iterations (i.e. the number of hash tables) and the minimum similarity threshold for merging spectra. To select the optimal default parameters, we evaluated the performance of msCRUSH using different parameter combinations ranging from 10 to 20 hash functions and from 25 to 150 iterations. The results from these experiments are shown in Supplementary Table S1 and S2. Overall, when 15-18 hash functions and 50-100 iterations were used, the clustering results are satisfactory in terms of purity, within-cluster entropy and running time. With the overall consideration of running time and cluster metrics, we selected 15 hash functions and 100 iterations as the default parameters in msCRUSH. We also evaluated msCRUSH using different minimum similarity thresholds, 0.5, 0.55, 0.6 and 0.65, respectively. As shown in the Supplementary Table S3, it appears different default similarity thresholds should be selected for clustering MS/MS spectra of different charges. Finally, we set the default values for the minimum similarity thresholds. Those thresholds are: 0.6 for spectra of charge 1+, 0.65 for spectra of charge 2+, 0.6 for spectra of charge 3+, and 0.55 for spectra of charge 4+. The optimal threshold may vary in different datasets. During the iterative process, msCRUSH uses a decaying function to gradually decrease the threshold from 0.9 to the respective minimum threshold. For the sake of consistency, we set 0.65 as the default value of minimum similarity threshold for both datasets of all charge states, then report clustering results of msCRUSH including purity, within-cluster entropy, within-peptide entropy and running time.

### Clustering Performance

We evaluated the performance of msCRUSH on two testing datasets in comparison with PRIDE Cluster, the state-of-the-art spectra clustering algorithm that has been applied to the massive MS/MS spectra in the PRIDE data repository.^29^ The results are summarized in Table 1. Overall, msCRUSH runs 3.2-12.1 times faster than PRIDE Cluster, and the acceleration rate is higher for larger datasets. On the largest dataset tested so far, i.e., the B-2+ dataset containing 11.5 million of doubly charged spectra, msCRUSH completed in 783 seconds (≈13 minutes) over 100 iterations, 12.1 times faster than PRIDE Cluster that completed in 9,460 seconds (≈158 minutes), both using 40 threads. Note that due to the running time constraints, PRIDE Cluster only performed 4 iterations of clustering. As a result, PRIDE Cluster often generated incomplete clustering results, with typically more clusters than those reported by msCRUSH. For example, for the dataset B-2+, msCRUSH grouped the input 11.5 million spectra into 5.6 million clusters, while PRIDE Cluster reported 7.7 million clusters, 36.8% more than the msCRUSH results. This reduction in numbers of clusters (and the corresponding numbers of consensus spectra) can further speed up the subsequent step of the database searching for peptide identification, as we will discuss below.

Next, we evaluated the quality of clusters reported by msCRUSH and PRIDE Cluster using different evaluation metrics. To compute the evaluation metrics, we used the database searching results from MSGF+^10^ with the false discovery rate of 1% on peptides level for the original datasets with un-clustered MS/MS spectra. We removed the common singletons (i.e. clusters with only one spectrum) from clustering results of msCRUSH and PRIDE Cluster. As shown in Table 1, msCRUSH generated spectra clusters with high purity (close to 1), indicating the majority of spectra in each cluster are identified as the same peptides. PRIDE Cluster generated clusters with slightly higher purity values comparing to msCRUSH. The purity of both algorithms on the large dataset B-2+ differs only by 0.009. The results of the within-cluster entropy demonstrated the same trends, although the within-cluster entropy measures the diversity of peptides within one cluster that is reversely correlated with purity measure. Interestingly, the within-peptide entropy that measures the distribution of spectra of the same peptides in different clusters, demonstrated significant difference between msCRUSH and PRIDE Cluster: msCRUSH shows much lower within-peptide entropy than PRIDE Cluster, indicating that msCRUSH has grouped the spectra from the same peptides more effectively than PRIDE Cluster. Finally, it is worth noting, among the spectra clusters reported by both algorithms, the majority are singletons (see Supplementary Figure S1 for the size distribution of the clusters), which may represent the noise spectra acquired in the LC-MS/MS experiments. However, msCRUSH outputs fewer singleton clusters than PRIDE Cluster before removing common singletons (e.g. 5,012,516 vs. 7,001,404) on the B-2+ dataset, again indicating msCRUSH can effectively group similar spectra into the same cluster. We also evaluated the quality of clusters from PRIDE Cluster and msCRUSH after removing all singletons from the clustering results, and we observed similar results to those results on clusters removing only the common singletons (see Supplementary Table S4).

### Database searching using consensus spectra

One important application for spectra clustering is to reduce the number of spectra while improving their quality by using the consensus spectra after spectra clustering^12^ for peptide identification. To test the performance of this approach, we used peptide search engines MSGF+ and Mascot, respectively, to identify the consensus spectra, each representing a spectra cluster, reported by both msCRUSH and PRIDE Cluster, and compared them with the peptide identification results from the original MS/MS datasets using the same peptide search engines. As depicted in Figure 3(a) and 3(c), using consensus spectra generated by msCRUSH and PRIDE Cluster, respectively, can both significantly reduce the time for data searching through the MSGF+ search engine (see Supplementary Table S5 for similar results of Mascot), as the number of consensus spectra is significantly smaller than that of the input un-clustered MS/MS spectra. Note that the running time of both msCRUSH and PRIDE Cluster are much shorter than the time of database search using MSGF+ (e.g. it takes 13 and 158 minutes for msCRUSH and PRIDE Cluster, respectively, to cluster the 11.5 million spectra in the B-2+ dataset, while the database searching takes 4-5 thousands of minutes on consensus spectra reported by msCRUSH and PRIDE Cluster, respectively). For example, on the B-2+ dataset with 11.5 million spectra, using the consensus spectra from msCRUSH can speed up the database searching time by 2.37x (reduced from 9, 228 to 3, 894 minutes), compared to the 1.72x speedup by PRIDE Cluster (reduced from 9, 228 to 5, 370 minutes) when MSGF+ is used. Similar results (see Supplementary Table S5) have been obtained for Mascot, we can achieve a 2.23x (reduced from 654 to 293 minutes) speedup using the consensus spectra from msCRUSH, and an 1.68x (reduced from 654 to 391 minutes) speedup using the consensus spectra from PRIDE Cluster. The maximum speedup can be achieved on the A-2+ dataset (with 415k spectra) for both search engines: 4.26x (reduction from 301.1 to 70.7 minutes) for MSGF+, and 5.95x (reduction from 43.4 to 7.3 minutes) for Mascot.

**Figure 3:**
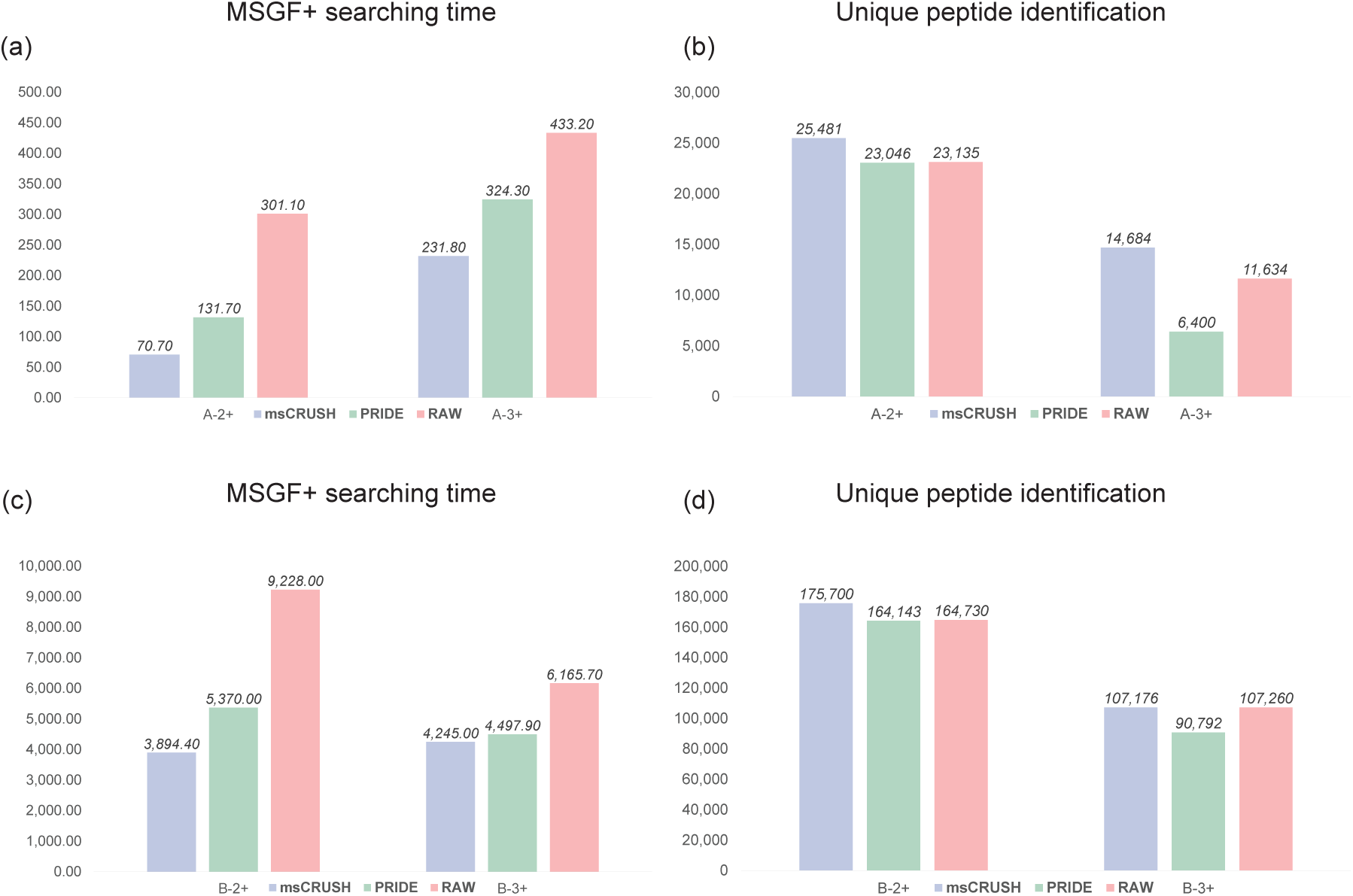
Comparison of the database searching time in minutes and unique peptide identification through the MSGF+ search engine using the raw un-clustered MS/MS spectra and the consensus spectra generated by msCRUSH and PRIDE Cluster, respectively, as input. (a) MSGF+ searching time on Dataset A; (b) Unique peptide identification on Dataset A; (c) MSGF+ searching time on Dataset B; (d) Unique peptide identification on Dataset B.

Using the consensus spectra can indeed improve the overall peptide identification results, and the improvement is strikingly pervasive across all charge states and search engines (either MSGF+ or Mascot). As shown in Figure 3(b) and 3(d), on almost all tested datasets, more unique peptides were identified through the MSGF+ search engine from the input of consensus spectra generated by msCRUSH than those identified from the input of original, un-clustered MS/MS spectra, when the same peptide-level false discovery rate (1%) is applied. Though for the B-1+ dataset, when Mascot was used, the number of identified unique peptides from consensus spectra produced by msCRUSH is less than that from the un-clusted, raw MS/MS spectra, but is still much more than the identification result from the input of consensus spectra reported by PRIDE Cluster (see Supplementary Figure S2). On average, 5% more unique peptides can be identified by using MSGF+ as the search engine from the consensus spectra generated by msCRUSH, while 4% such improvement when Mascot was used as the search engine. In an extreme case, on the dataset of A-3+ (of triply charged spectra), 26% more unique peptides can be identified (an increase from 11, 634 to 14,684) by MSGF+ from the input consensus spectra generated by msCRUSH. It is worthy noting, different cosine similarity thresholds for merging similar spectra, such as 0.5, 0.55, 0.6 and 0.65, lead to similar number of identified unique peptides from consensus spectra produced by msCRUSH when tested on all datasets, as depicted in Supplementary Table S6. In comparison, using the consensus spectra generated by PRIDE Cluster often leads to the reduced number of identified unique peptides (see Figure 3 and Supplementary Figure S2 and S3). For instance, on dataset A-3+, 50% less unique peptides were identified (a decrease from 11,634 to 6,400) by MSGF+ from the input of consensus spectra reported by PRIDE Cluster. This observation implies that the high quality of spectra clustering achieved by msCRUSH can improve the quality of the consensus spectra generated from the clusters, which will then improve the peptide identification results using these consensus spectra.

Finally, we investigated the difference among the identified peptides from the input of the un-clustered spectra, and the consensus spectra generated by msCRUSH and PRIDE Cluster, respectively. Figure 4 demonstrates that on each of the datasets A-2+, A-3+, B-2+ and B-3+, the number of unique peptides identified on the un-clustered, raw spectra overlaps significantly with the unique peptides identified from the consensus spectra generated by msCRUSH and those by PRIDE Cluster. On all datasets (see Supplementary Figure S2 and S3), the results from the input of the consensus spectra generated by msCRUSH share more identified unique peptides with the results from the input of un-clustered original spectra, than the results from the input of consensus spectra generated by PRIDE Cluster, regardless of which peptide search engine (MSGF+ or Mascot) is used. This result again showed that msCRUSH algorithm can produce accurate consensus spectra representing most spectra in the original datasets.

**Figure 4:**
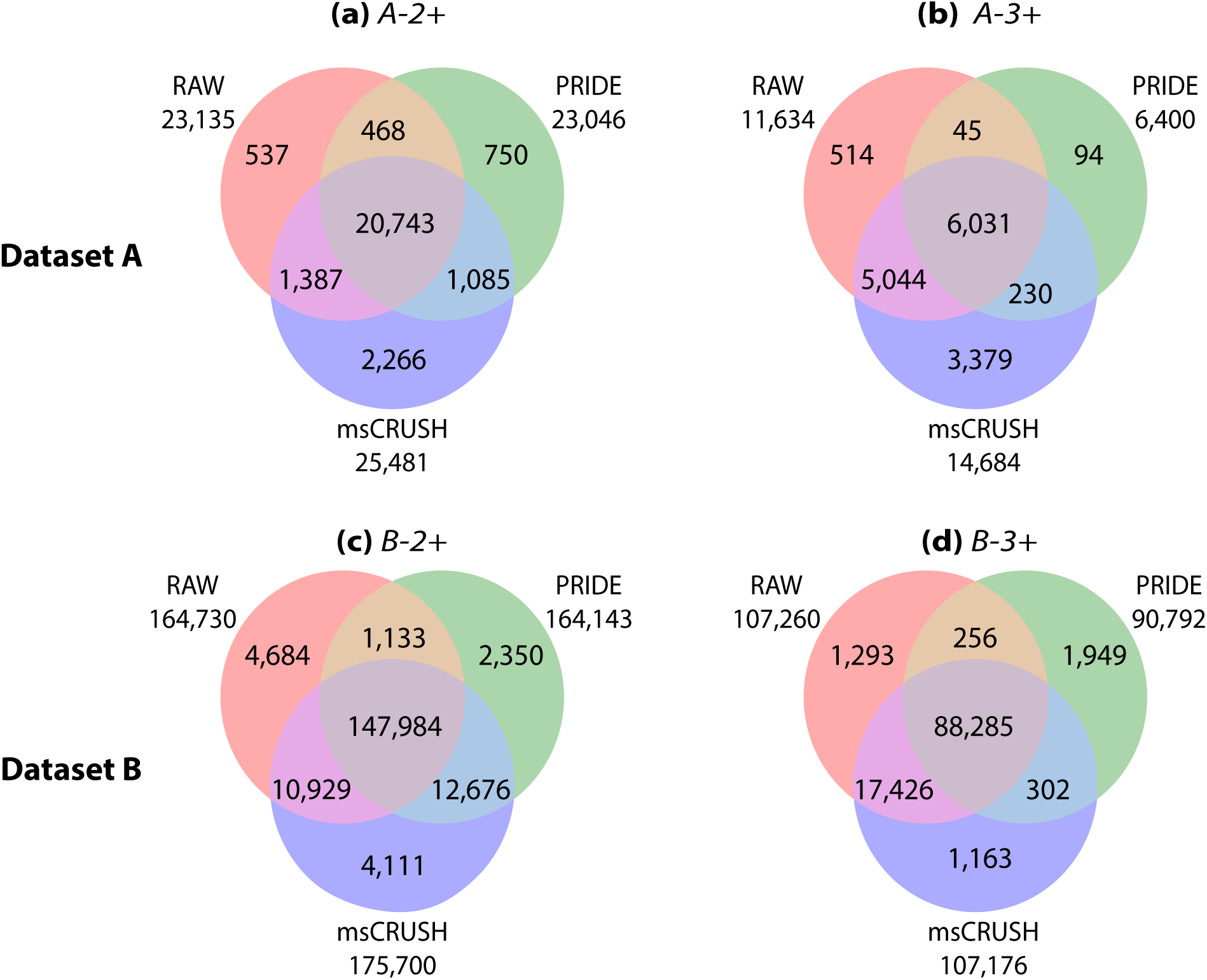
Venn diagrams demonstrating the overlap of the identified unique peptides using consensus spectra (i.e., generated by PRIDE Cluster and msCRUSH, respectively) and the original spectra in dataset A, B with charge 2+ and charge 3+, respectively, as input. MSGF+ was used for database searching. (a) Unique peptides identified from dataset A-2+ (b) Unique peptides identified from dataset A-3+ (c) Unique peptides identified from dataset B-2+ (d) Unique peptides identified from dataset B-3+.

## Discussion

We proposed a rapid algorithm msCRUSH for tandem mass spectra clustering with high sensitivity and accuracy. msCRUSH exploits the locality sensitive hashing (LSH) technique to speed up the similarity comparison between tandem mass spectra. To alleviate the approximation incurred by LSH, which may miss some similar spectra, an iterative clustering strategy is adopted: with more iterations, similar spectra are more likely to be grouped into a same cluster. Fewer iterations of clustering can lead to further speedup but may reduce sensitivity of spectra clustering. In fact, the current spectra clustering algorithms such as PRIDE Cluster typically applied as few as four iterations because of the constraints of running time. By using LSH, msCRUSH can perform much more iterations (100 by default), generating fewer and larger clusters, as well as more accurate consensus spectra, which can improve the peptide identification results when using the consensus spectra as input for database searching. Notably, these improvements were achieved simultaneously with a 3-12 folds of speedup comparing to PRIDE Cluster, specifically on the large dataset of 18.4 million spectra.

Our evaluation showed that spectra clustering can improve peptide identification because the consensus spectra generated from spectra clusters exhibit higher quality than the original spectra. Nevertheless, many consensus spectra were not identified even though they represent large clusters of spectra. For example, in the largest testing dataset B-2+, a total of 89,744 consensus spectra were generated by msCRUSH algorithm from clusters containing 10 or more MS/MS spectra; among them, 38,246 (42.6%) consensus spectra were not identified by either MSGF+ or Mascot through the database searching against the Uniprot human protein database. Many of these spectra perhaps result from human peptides containing mutations or post-translational modifications (PTMs), and thus are worth further investigation. When applied to clinical proteomic datasets (e.g., acquired from samples of case/control subjects), spectra clustering algorithms enable a spectrum-centric approach to quantitative biomarker discovery, which can focus on the interpretation of the clusters of spectra corresponding to parent ions with differential abundances in the case and control samples. For this approach to succeed, however, it is crucial to group all spectra from the same peptides into the same clusters; otherwise, the quantitative comparison becomes impossible. The high sensitivity of msCRUSH clustering algorithm makes it particularly applicable for this approach.

Although it is designed to cluster MS/MS spectra of peptides acquired from proteomic experiments, msCRUSH can be directly applied to clustering the MS/MS spectra of other molecules, in particular metabolites from metabolomics.^36^ The enhanced spectrum quality after spectra clustering may also improve the identification of MS/MS spectra in these applications.

## Acknowledgement.

This work was supported by the NIH grant 1R01AI108888 and the Indiana University (IU) Precision Health Initiative (PHI). We thank helpful discussions with Haoyu Zhang and Dr.Qin Zhang at Indiana University, Bloomington. We are also grateful to Dr.Yongan Zhao and Dr. Xiao Wang (University of Maryland, College Park), who provided helpful discussions about implementation of multithreading.

